# A Newly Identified Gene Controls Lignin Acetylation in Poplar

**DOI:** 10.1101/2025.10.06.680708

**Authors:** Nuoendagula, Fachuang Lu, John Ralph

## Abstract

Acetylation in plant cell walls, including lignin, plays a critical role in biomass processability. High acetate content facilitates uncatalyzed, additive-free hot-water or steam hydrolysis, releasing acetic acid without the need for mineral acids or other chemical pretreatments. Despite the prevalence of acetylation in certain plant species, the genes and enzymes responsible for lignin and xylem acetylation remain largely uncharacterized. Here, we report the identification and functional characterization of an acetyl-CoA:monolignol transferase gene in Populus. Overexpression of this gene in transgenic poplars led to elevated acetate levels in lignin without negatively impacting plant growth or development. Acetate accumulation was positively correlated with gene expression, and among 18 independent transgenic lines, several showed at least a sevenfold increase compared to wild-type controls. This work establishes a genetic basis for lignin acetylation and provides a promising strategy to engineer biomass with improved pretreatability, enhancing the efficiency of biofuel and bioproduct production.

## Introduction

Lignin, a complex heteropolymer, holds significant intrigue due to its structural complexity, multifaceted roles in plant defense, and barrier properties that impede the enzymatic breakdown of polysaccharides into fermentable sugars. Despite its preeminence as a sustainable source of aromatic compounds, lignin remains underutilized.

Lignin is synthesized through the oxidative coupling of various phenolic compounds, deriving primarily from the monolignols, a series of variously methoxylated hydroxcinnamyl alcohols: *p*-coumaryl alcohol, the *p*-hydroxyphenyl (H) unit precursor, coniferyl alcohol, the guaiacyl (G) unit precursor, and sinapyl alcohol, the syringyl (S) unit precursor (Freudenberg et al., 1968). It is increasingly realized that numerous other phenolic monomers may be involved in lignification. These include various monolignol conjugates (Ralph et al., 2023). Monolignols and their conjugates originate from both primary and secondary metabolic pathways in plants, with the shikimate-phenylpropanoid pathway playing a central role. Various unusual lignin monomers, such as flavonoids and other polyphenols, have also been implicated in lignification (Lan et al., 2015; del Río et al., 2017; del Río et al., 2018; del Río et al., 2022; Ralph et al., 2023). It is widely accepted that, primarily due to differences in oxidation potential and the relative stabilities of the produced radicals, the G- and S-units constitute the primary backbone of lignin. Minor H-units and other structural components are predominantly located at the terminal ends of the lignin polymer (Lapierre et al., 1988a; Lapierre et al., 1988b; Takahama et al., 1996; Kobayashi et al., 2005; Hatfield et al., 2008).

The β–O–4 bond in arylglycerol-β-aryl ether units, the most prevalent inter-unit linkage in lignin, accounts for approximately 50–60% of the lignin structure in both softwoods and hardwoods. Other lignin dimeric units, including the resinol (β–β), phenylcoumaran (β–5), biphenyl (5–5’), dibenzodioxocin (5– 5’/4–O–β), and spirodienone (β–1) units, have been identified in natural lignins (Karhunen et al., 1995; Katahira et al., 2018; Ahmad et al., 2020; Matsushita et al., 2020; Ralph et al., 2023). Benzodioxanes (β– O–4, albeit in cyclic structures due to the internal trapping of the quinone methide intermediate) emerge when OMT-deficient plants incorporate caffeyl alcohol or 5-hydroxyconiferyl alcohol into the monomer pool (Marita et al., 2001; Ralph et al., 2001; Marita et al., 2003; Jouanin et al., 2004; Morreel et al., 2004; Lu et al., 2010; Wagner et al., 2011; Chen et al., 2012; Ando et al., 2021). The linkage type, monomer composition, and polymerization degree of lignin exhibit variations not only across different plant species but also among distinct cell types, tissue types, and developmental stages within the plant (Chen et al., 2012; Li et al., 2016; Blaschek et al., 2020; Ménard et al., 2022). These disparities have become a primary focus of research in the areas of deconstruction, engineering, and utilization of lignin.

Recent research has provided valuable insights into the deconstruction, engineering, and utilization of lignin polymers. For instance, CRISPR/Cas9-mediated downregulation of the monolignol biosynthetic gene *Cinnamoyl-CoA Reductase* (*CCR*) in *Populus* has been demonstrated to enhance saccharification efficiency by up to 41%, depending on the type of pretreatment applied (De Meester et al., 2020). Natural mutants of *Cinnamyl Alcohol Dehydrogenase* (*CAD*) in *Morus* and pine, as well as *CAD*-RNAi-Poplar, have exhibited enhanced chemical pulping and saccharification performance (Sederoff et al., 1986; Ralph et al., 1997; Van Acker et al., 2017; Ikeda et al., 2021). *O*-Methyltransferase-deficient plants produce regular linear lignins; C-lignin, a homopolymer of caffeyl alcohol, can be efficiently depolymerized through a hydrogenolytic method, yielding pure catechol monomers with a yield of up to 90% (Li et al., 2018).

Lignins enriched with monolignol ester conjugates, commonly referred to as monolignol conjugates, have garnered significant attention due to their chemically labile ester linkages and their potential for diverse industrial applications (Bartley et al., 2013; Wilkerson et al., 2014; Timokhin et al., 2020; Smith et al., 2022a; Smith et al., 2022b; Ralph et al., 2023; Karlen et al., 2024; Unda et al., 2024). Monolignol ferulate conjugates are particularly noteworthy because they induce the formation of “zip-lignins” in plants due to their propensity to incorporate into the lignin polymer backbone because of their reactive phenolic groups at both ends (Wilkerson et al., 2014; Ralph et al., 2019). Lignins incorporating monolignol conjugates have garnered industrial interest primarily stemming from their relative ease of deconstruction (Wilkerson et al., 2014; Kim et al., 2017; Zhou et al., 2017). Although the molecular mechanisms underlying the acylation and incorporation of acyl moieties into lignin in lignified plant cells remain elusive, genes responsible for these monolignol conjugates belong to the plant-specific BAHD acyltransferase family (D’Auria, 2006). These genes have been identified in numerous plant species, including genetically modified plants that have undergone modification in their lignin and related biosynthetic pathways. Notable examples include feruloyl-CoA:monolignol transferase (FMT) (Wilkerson et al., 2014), *p*-coumaroyl-CoA:monolignol transferase (PMT) (Marita et al., 2014; Petrik et al., 2014; Smith et al., 2022a), *p*-hydroxybenzoyl-CoA:monolignol transferase (*p*HBMT) (de Vries et al., 2022), and benzoyl-CoA:monolignol transferase (BMT) (Chedgy et al., 2015; Kim et al., 2020). Research indicates that these monolignol transferase genes are present in a diverse range of plant species (Karlen et al., 2016; Kim et al., 2020). Several other studies have demonstrated that engineering these enzymes into crop plants enhances deconstruction and saccharification efficiency (Grabber et al., 2008; Grabber et al., 2012; Tobimatsu et al., 2012; Ralph et al., 2023). Several atypical conjugates have also been detected in the lignin polymer, including tyramine ferulate in both native and transgenic tobacco (Ralph et al., 1998a), for which the responsible biosynthetic gene, well-known in tobacco and potato (Schmidt et al., 1999), has been identified (Farmer et al., 1999), and rosmarinic acid, a caffeate conjugate found in Lamiaceae and various other plant families (Petersen et al., 2003; Sander et al., 2011; Tobimatsu et al., 2012).

Although acetate-monolignol conjugates are implicated in numerous plant species, the BAHD gene specifically responsible for the acetyl-CoA:monolignol transferase (AMT) activity remains unreported to date. For example, γ-acetylated sinapyl alcohol was initially identified as the precursor to lignins acylated at their γ-hydroxy positions in kenaf bast fibers; the identification of β–β-coupled units that were tetrahydrofurans and not the usual resinols could only arise during lignification with pre-acylated monolignols (del Río et al., 2007; Lu et al., 2008). However, an attempt to deduce the gene instead led to an unexpected *p*-coumaroyl-CoA:monolignol transferase (Mottiar et al., 2023). In this study, a poplar acetyl-CoA:monolignol transferase gene (*PtAMT1*) [Potri.007G003800 (1479bp)] was screened. Its overexpression, driven by the lignified cell-specific Arabidopsis cinnamate-4-hydroxylase (*C4H*) gene promoter, in hybrid aspen (*Populus tremula* × *Populus alba* clone INRA 717-1B4) resulted in elevated acetate levels in lignin without compromising plant growth. Our findings demonstrate that the AMT enzymes identified herein exhibit high specificity for acetyl-CoA:monolignol transferase activity, thereby facilitating the targeted modulation of lignin biosynthesis of plants for practical applications.

## Results and Discussion

### Identification of the Poplar putative *PtAMT1* gene and phylogenetic analyses

*PtAMT1* was identified through homologous alignment with other BAHD acyltransferase genes in a database. Well-characterized monolignol acyltransferase genes were excluded using multiple sequence alignment and phylogenetic analysis that incorporated both distance-based and likelihood-based methods. Unique mutations or structural features in the target gene were then identified. As depicted in Supplemental Fig. S1B, *PtAMT1* (an AMT, highlighted) clustered into the same clade as the coniferyl alcohol acyltransferase gene (*CFAT*) that plays an important role in the production of isoeugenol by catalyzing the initial conversion of coniferyl alcohol to coniferyl acetate (Dexter et al., 2007; Zhang et al., 2019; Wang et al., 2023). The 1479 bp coding DNA sequence (CDS) of *PtAMT1* encodes a polypeptide of 492 amino acids with a predicted molecular mass of 53.99 kDa. The *PtAMT1* gene retained the BAHD acyltransferase conserved residues -HXXXD- and -DFGWG- and had 56–70% amino acid identity compared to coniferyl alcohol acyltransferase (*CFAT*) (Supplemental Table S1). However, the protein exhibited a maximum of 27% amino acid identity with hydroxycinnamoyl-CoA:shikimate hydroxycinnamoyltransferase (HCT), a key enzyme that acylates shikimate with *p*-coumarate (via its CoA thioester) on the pathway to monolignols, and a maximum of 34% amino acid identity with proteins from genes encoding monolignol transferases such as *FMT*, *PMT*, *pHBMT*, and other genes involved in cell wall acylation.

The *PtAMT1* gene identified in this study is present as a single copy in the poplar genome and has not undergone whole-genome duplication. Transcriptome data from Phytozome/Department of Energy’s Joint Genome Institute (JGI) indicate low expression levels of this gene in primary tissues, such as leaves, roots, and stems, suggesting its minimal activity in these organs. However, detailed transcriptomic analyses have revealed its potential involvement in acyl-lipid and phenylpropanoid metabolism, particularly within cork tissues (Rains et al., 2018). Comparative sequence analysis highlights *PtAMT1* as having the highest sequence identity with known hydroxycinnamoyl-CoA:monolignol and (hydroxy)benzoyl-CoA:monolignol transferase genes. *PtAMT1* exhibits a 69.87% sequence identity with the cinnamyl alcohol:acetyl-CoA transferase (*CAAT1*) gene from *Larrea tridentata*, the expressed protein of which demonstrates the highest substrate specificity for *p*-coumaryl alcohol, followed by caffeyl alcohol, coniferyl alcohol, and relatively lower specificity for sinapyl alcohol and 5-hydroxyconiferyl alcohol (Kim et al., 2014). In contrast, *PtAMT1* shares a lower sequence identity (56.12%) with the coniferyl alcohol:acetyl-CoA transferase (*CFAT* gene) from *Petunia hybrida*, the protein of which exhibits the highest substrate specificity for coniferyl alcohol compared to other analogs (Dexter et al., 2007). Note that *CFAT* and other monolignol-specific genes may all be regarded more generically as *CAAT* genes. *PtAMT1* also exhibits higher (61%-67%) sequence identity with several other *CFAT/CAAT* candidate genes that have been characterized and analyzed in various plant species with the objective of identifying genes involved in the production of valuable chemicals, such as eugenol or isoeugenol. For example, transgenic *Petunia* plants expressing a *CFAT*-RNAi construct exhibited decreased levels of isoeugenol (Dexter et al., 2007). Additionally, CFAT in *Prunus mume* is localized in both the cytoplasm and nucleus (Zhang et al., 2019), and an acetyl-CoA:sinapyl alcohol transferase has been identified in *Euphorbia lathyrism* (Wang et al., 2023). All these proteins possess the conserved HXXXD motif that has been established to be involved in the catalytic activity of BAHD enzymes (Bayer et al., 2004; Kruse et al., 2023). However, the DFGWG motif is absent in some of these proteins and, particularly in the CFAT from *Petunia hybrida*, as are DFGWG variants. This observation may support the hypothesis that the DFGWG motif does not directly contribute to substrate binding or catalysis (Galaz et al., 2013). Notably, overexpression of *CFAT* from *Petunia hybrida* in *Arabidopsis* plants, driven by the same promoter used in this study, resulted in no acetylation of lignin in the transgenic plants (data not shown). This outcome could be attributed to differences in metabolic flux between *Arabidopsis* and *Populus* but may also suggest a potential role for the DFGWG motif in these proteins, as well as possible differences in the catalytic sites or solvent channels of the enzyme (Galaz et al., 2013), potentially driven by the highly divergent evolution of the BAHD gene family (St-Pierre et al., 2000). We anticipate conducting further analysis of the active sites of these enzymes and their interaction patterns with substrates, along with both *in vitro* and *in planta* validation of these genes.

### Validation of PtAMT1 *in planta*

To assess the acetyl-CoA:monolignol transferase activity of the PtAMT1 enzyme *in planta*, the coding sequence of *PtAMT1* was transformed into the model tree hybrid aspen, *Populus tremula* × *Populus alba* clone INRA 717-1B4, under the control of a strong lignin-targeting cell-specific promoter, the *Arabidopsis* cinnamate-4-hydroxylase (*AtC4H*) promoter (Bell-Lelong et al., 1997). Eighteen transgenic lines were selected using genomic PCR, and three transgenic poplars with the highest, lowest, and medium levels of the target gene’s expression were selected based on semi-quantitative RT-PCR analysis (Supplemental Fig. S1A). The transgenic plants exhibited normal growth, indicating that at least a 7-fold increase in acetate level on lignin does not compromise the poplar cell wall architecture or plant growth and development. We have not yet been able to locate *PtAMT1* gene homologs in the background poplar genome in an existing database. Trace amounts of gene expression were observed in the cDNA from wild-type poplar (Supplemental Fig. S1A), suggesting that the background poplar expresses a gene similar to the target gene, but at a low level.

### Authenticating lignin acetylation via DFRC

Lignin acylation was initially regiochemically contentious (Nakamura et al., 1976) until NMR methods unequivocally established that *p*-coumarate in grass lignins exclusively attaches to the γ-OH of lignin sidechains (Ralph et al., 1994). The DFRC method (Lu et al., 1997) cleaves the predominant arylglycerol-β-aryl ether (β–O–4) bonds in lignin while preserving esters and amides, a discovery of considerable practical value (Lu et al., 1999, 2014; Lu et al., 2015; Regner et al., 2018; del Río et al., 2020; Ralph et al., 2023; del Río et al., 2025). This method, in conjunction with NMR methods (subsequently discussed), has been pivotal in establishing that lignin acylation occurs through the incorporation of acylated monolignols during polymerization (Ralph, 1996). For instance, *p*-coumaroylated monomers are released from grasses (Lu et al., 1999; Petrik et al., 2014) and ‘all’ commelinid monocots (Regner et al., 2018), and *p*-hydroxybenzoylated monomers are released from certain angiosperms such as poplar (Lu et al., 2015; Regner et al., 2018) that have long been recognized as possessing lignins acylated with *p*-hydroxybenzoate (Smith, 1955).

The establishment of relative levels of γ-acetylation was achieved through a modification of the DFRC method in which its acetate-based reagents are replaced by their propionate analogs in the so-termed DFRC′ method (Ralph et al., 1998b; Lu et al., 2008). Lignin acetylation levels from released monolignol γ-acetates have been reported for various angiosperms (del Río et al., 2007). For instance, in sisal, 78% of G-units and 50% of S-units are acetylated; in kenaf, 59% of S-units and 9% of G-units are acetylated (del Río et al., 2007). Lignin acetylation has not been reported in Gymnosperms (del Río et al., 2007).

To assess the impact of the *PtAMT* gene on lignin in our poplar transgenics, we employed the DFRC′ method to identify natural acetates on lignin units, Fig. 1. Utilizing extract-free cell wall residue samples, our DFRC′ results demonstrated the release of guaiacyl and syringyl monolignol acetate conjugates **G**^Ac^ and **S**^Ac^ from the transgenic poplar lines, confirming that the *PtAMT* gene we selected possesses acetyl-CoA:monolignol transferase activity *in planta*. As depicted in Fig. 1C, transgenic poplar with the highest expression level exhibited 17.4% acetylated **S** monomer **S**^Ac^, 21.8% acetylated **G** monomer **G**^Ac^, and a total of 18.7% acetylated monomers, compared to 3.3%, 0.8%, and 2.5% in wild-type poplar. This suggests that *PtAMT1* encodes an enzyme that preferentially catalyzes G-monomer acetylation. The S/G ratio in some of the transgenic plant lines was slightly lower than in wild-type poplar (Fig. 1B). The total lignin measured using the thioglycolic acid method remained unchanged, indicating that the level of lignin acetylation in the selected transgenic plants did not significantly alter the lignification process.

**Figure 1.**
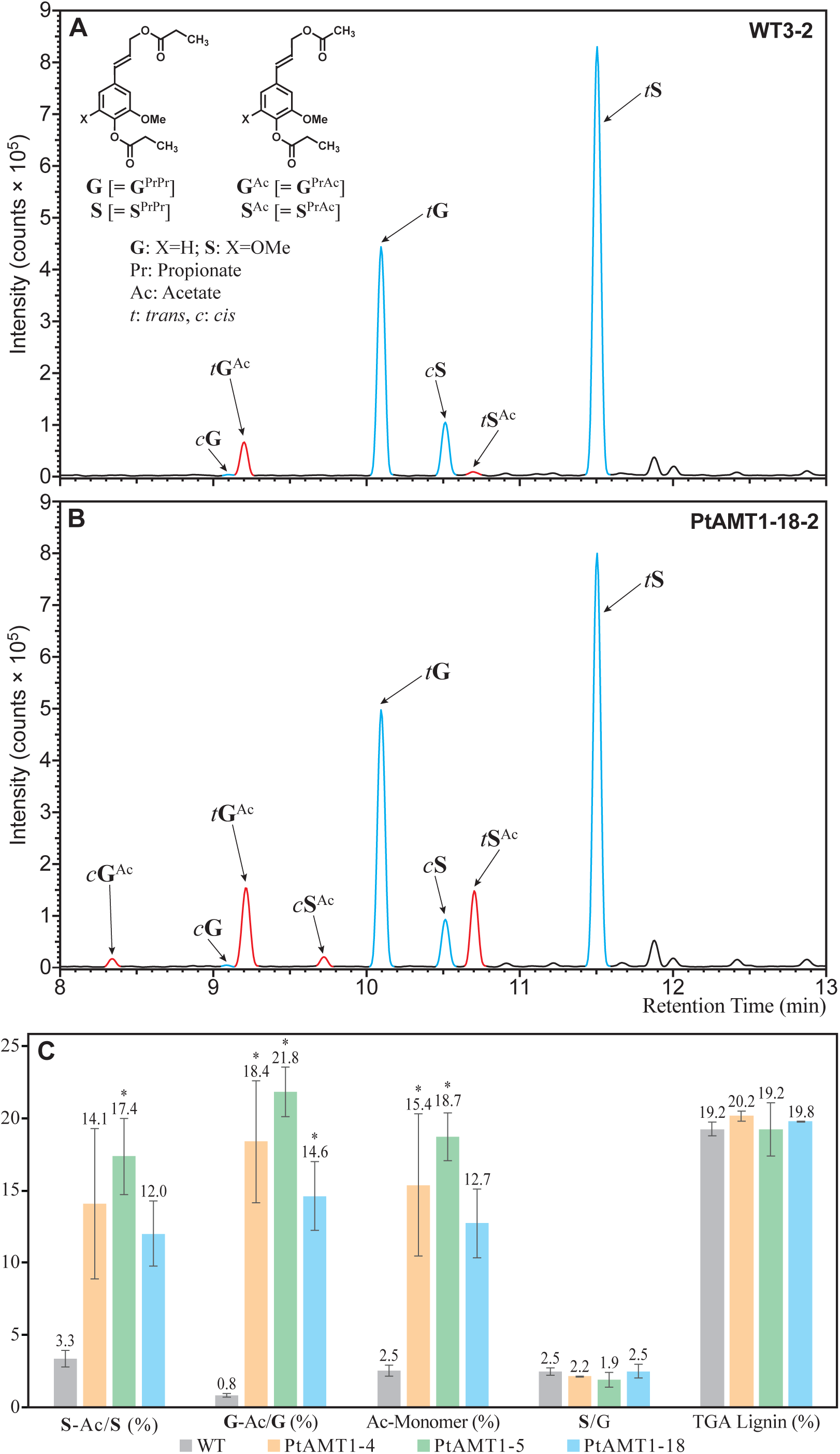
Monolignol acetate conjugates by DFRC′ analyses. **A)** WT poplar extracted-ion chromatogram of GC-MRM-MS transitions monitored for DFRC products. **B)** Analogous spectra from a PtAMT1 transgenic line. **C)** The percentage of various DFRC-releasable monolignol acetate units obtained by GC-MRM-MS, along with the S/G determination from the monomer, and the total lignin contents determined by the thioglycolic acid method. Data are means of triplicate or duplicate analyses. Asterisks indicate statistical significance.

### Delineating the structural aspects of lignin acetylation through NMR analysis

To elucidate the impact of the *PtAMT1* gene on lignin structure and its constituent units (with their characteristic inter-unit linkages), various 2D homo- and heteronuclear NMR experiments were conducted, primarily 2D-HSQC, complemented by the powerful and insightful 3D-TOCSY-HSQC experiment. These experiments were supported by model compounds for assignment authentication. Fig. 2A illustrates the oxygenated-aliphatic region of the HSQC spectrum from an enzyme lignin (EL) isolated from the *PtAMT1* transgenic. In comparison to the WT control (Fig. 2B), several novel lignin-derived peaks are discernible. These peaks, in comparison with model data (Supplemental Fig. S2) and 3D (Supplemental Figures S3 and S4) provide diagnostic evidence for the acetylation of the γ-hydroxy group of β-ether **A**, phenylcoumaran **B**, and cinnamyl alcohol end-group **F** units. Although such qualitative HSQC spectra lack quantitative precision, the volume-integration of representative α-H/C correlation contours facilitates the estimation of relative levels and highlights the alterations in interunit linkage distribution (Mansfield et al., 2012). Relative levels of aromatic units were also measured from integrating proton-carbon 2-correlations for **G** and 2/6-correlations from **S** and ***p*HB** units that are not within the plotted region but are summarized in the distribution table in the bottom-right of the spectra in Fig. 2 and are illustrated and detailed in Supplemental Fig. S5.

**Figure 2.**
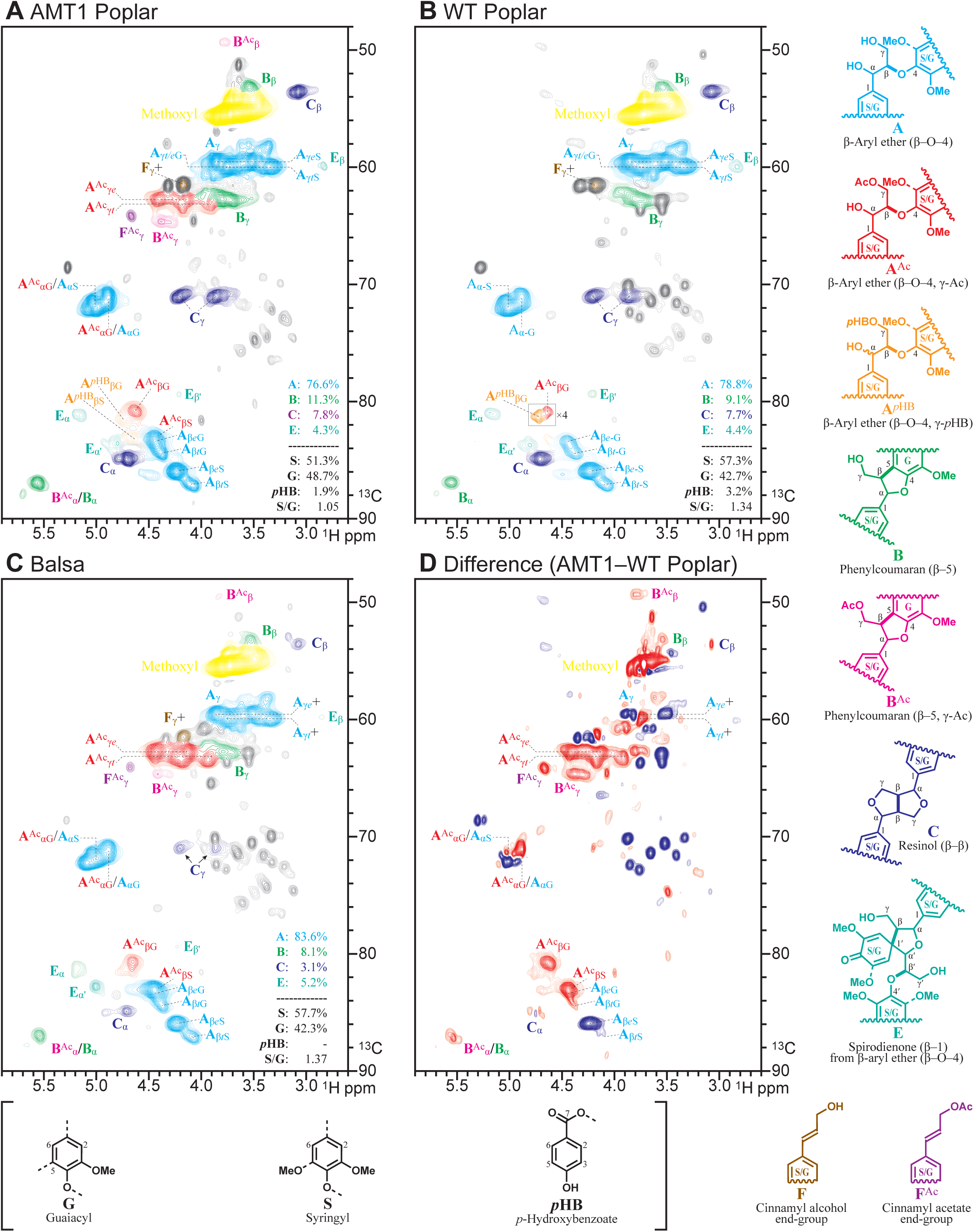
Structural/compositional profiling of enzyme lignins (EL) by 2D-^1^H–^13^C-correlation NMR spectroscopic analysis – partial HSQC spectra (the oxygenated aliphatic region for profiling of the interunit linkages. **A)** The *PtAMT1* transgenic with the acetyl-CoA:monolignol transferase expressed. **B)** The WT control poplar. **C)** Balsa that, as revealed here, natively has an analogous lignin acetylation profile. **D)** The 2D difference spectrum (AMT1 minus WT) using the best nulling of the **S**_2/6_ signal, revealing the compositional differences. Warm colors represent components elevated in the transgenic line compared to WT (subject to this **S**_2/6_ signal nulling), whereas cold colors represent depleted components relative to the syringyl peaks. As readily noted, peaks from γ-acetylated components were markedly differential and in fact were scarcely present in the WT even when the spectra are viewed down toward the baseplane noise level. The main correlation peaks are color-coded to match the structures shown. Note that we use the standard designation of lignin units characterized by their interunit linkages, **A** for β-ethers (β–O–4), **B** for phenylcoumarans (β–5), **C** for resinols (β–β), and **E** for spirodienones (β–1/β–O–4); structures **D** for dibenzodioxocins (5–5) were not evident here and are therefore not labeled and the structure is not shown. The lighter (40%) contours are from a 2-fold intensity expansion to facilitate the identification of more minor peaks. The relative levels of key components were assessed by volume-integration, as summarized in the table in the lower-right corner of each figure.

In comparison to wild-type poplar, *PtAMT1* poplar exhibited a lower S/G ratio, consistent with the DFRC measurements from the monomers released by cleaving β-ethers (Figs. 2A-B, Supplemental Fig. S5). The *p*-hydroxybenzoate ***p*HB** level was slightly decreased in the transgenic, further corroborating the DFRC results. Among the primary characteristic units in the lignin, the relative level of β-ether units **A** was slightly lower and the level of phenylcoumarans **B** was slightly higher, both features primarily attributable to a decrease in the proportion of S-units in the transgenic poplar (Stewart et al., 2009; Samuel et al., 2014).

The lignin of the *PtAMT1*-upregulated poplar exhibited a comparable acetylation pattern to that of Balsa (Fig. 2C), which belongs to the same clade as poplar but is part of the Malvaceae family. Balsa lignin possesses a similar S/G ratio to the wild-type Poplar plant but lacks *p*-hydroxybenzoates that are characteristic of *Salix*, *Populus*, and *Palmae* species (Smith, 1955; Nakano et al., 1961; Landucci et al., 1992; Sun et al., 1999; Meyermans et al., 2000; Li et al., 2001; Lu et al., 2004; Rencoret et al., 2013; Lu et al., 2015; Karlen et al., 2017; Rencoret et al., 2018; Goacher et al., 2021). Notably, *p*-hydroxybenzoylation was not detected in the NMR spectrum of the highly acetylated S-rich lignins from kenaf, a member of the Malvaceae family (Kim et al., 2010; Mottiar et al., 2023). Although the molecular mechanisms underlying the disparities in lignin linkage distributions and acylation patterns necessitate further investigation, these findings, in conjunction with the observed acylation patterns in grasses, suggest that plants within the same family exhibit analogous acylation patterns.

A difference spectrum of the transgenic *PtAMT1* poplar minus the wild-type poplar lignin (Fig. 2D), obtained by nulling the **S**_2/6_ signal, delineates peaks from γ-acetylated components that are substantially increased in the transgenic (red-colored contours in this difference spectrum). Signals corresponding to β-H/C correlations from syringyl β-ether units (**A**_βS_) are displayed in cold colors, indicating their depleted levels, whereas the peaks from the acetylated components, including **A**^Ac^_βS_ and **A**^Ac^_βG_ (along with guaiacyl β-ethers **A**_βG_), displayed in hot colors, are all relatively elevated. This observation aligns with the DFRC results, suggesting a lower abundance of syringyl β-ether units in the transgenic.

Other notable differences in the transgenic include the distinct γ-acetylated phenylcoumaran **B**^Ac^ peaks for **B**^Ac^_β_ and **B**^Ac^γ ; the **B**^Ac^_α_ peak is not resolved from the normal **B** peak. Additionally, although the normal γ-peak **F**_γ_ from cinnamyl alcohol end-units overlaps with contaminating protein peaks, its acetylated variant **F**^Ac^_γ_ emerges as a new discrete peak that is also elevated in the difference spectrum, Fig. 2D. All these correlation peaks, reliably authenticated from model data/spectra and from 3D NMR (see below, and in the Supplemental Materials), unequivocally demonstrate the substantial acetylation of the γ-hydroxyl in the transgenic’s lignin.

Although the peak dispersion is insufficient to enable relative quantification of all regions of these spectra through 2D-integration of the various peaks, certain relevant structures support the claim of a higher level of relative acetylation between the *PtAMT1* Poplar and the WT. For instance, the phenylcoumaran **B**^Ac^_β_ and **B**^Ac^_γ_ peaks are well-dispersed and well-resolved. In this case, the relative 2D integrals for the transgenic strain over the WT are at least 14-fold higher. We therefore hypothesize that acetate levels may exceed the 7-fold higher levels suggested by the DFRC data.

### Enhanced level of detail from 3D NMR coupled with model compound data

Supplemental Fig. S2 presents a parallel plot to that in Fig. 2A, incorporating pertinent model data that corroborates the accuracy of various assignments and expands upon them. For instance, it provides additional detail regarding the resolved isomers and delineates the peaks **A***^p^*^HB^_β_, analogous to those from **A**^Ac^_β_, from the β-ether units naturally acylated with *p*-hydroxybenzoate. These latter peaks are also discernable in the elevated (4×) detail in the WT poplar in Fig. 2B. Due to the overwhelming congestion and overlap, the isomers are not fully delineated on the plot. However, a few are constructed for the **A**_β_ correlations, labeled for **A**_β*e*G_, **A**_β*t*G_, **A**_β*e*S_, **A**_β*t*S_, in which *t* and *e* delineate the resolved *threo* and *erythro* isomers. All the data associated with these models will be deposited in this year’s release of our NMR Database (Ralph et al., 2025).

Of exceptional assignment value are the data from a 3D TOCSY-HSQC experiment. In this experiment, protons initially share their coupling-network-connections via TOCSY transfer, and the connections to the attached carbons from these protons are subsequently established by the HSQC sequence. As a result, ^1^H–^13^C-type HSQC spectra are unveiled at specific proton chemical shifts (Supplemental Fig. S3) or ^1^H–^1^H TOCSY spectra are revealed at given carbon shifts (Supplemental Fig. S4). The relative simplicity of these spectra, each showing only a limited number of correlation peaks, belies the remarkable power of such spectra to delineate detailed structural features involving specific units in lignin.

Supplemental Fig. S3 presents the F2-F3 spectra that are HSQC spectra at a given proton shift. Each is shown with the 3D plot in bold colors overlaid on the original 2D HSQC spectrum from Fig. S2A reproduced with a reduced opacity to highlight the emergence of the new correlations in these simple, structure-specific planes. Supplemental Fig. S3B, for instance, is taken at a proton shift of 5.59 ppm, primarily originating from the phenylcoumaran’s **B**_α_ proton. The observed correlations (in green) correspond to the H/C correlations for **B**_α_, **B**_β_, and **B**_γ_, confirming the assignments provided in Fig. 2A, Supplemental Fig. S2, and Supplemental Fig. S3A spectra. At a slightly lower proton chemical shift, at 5.51 ppm, correlations with the normal phenylcoumaran units **B** persist, but new correlations, in magenta, appear for **B**^Ac^_β_, **B**^Ac^_β_, and **B**^Ac^_γ_, further validating these assignments, particularly for the highly shifted and unique **B**^Ac^_β_ correlation. Most of the spectra, Supplemental Figs. S3D-M, demonstrate the various S and G β-ether correlations for non-acetylated and acetylated units. Notably, these spectra often separate the various *threo* and *erythro* isomers, a detail that is challenging to discern from the 2D HSQC spectrum alone. Further elaboration of this aspect can be found in the Supplemental Data. Finally, Supplemental Fig. S3N serves as evidence that other assignments can also be elegantly and unequivocally authenticated in such 3D NMR data. The plane through proton **C**_β_ at 3.04 ppm presents the set of resinol sidechain correlations, cleanly to carbons **C**_α_, **C**_β_, and **C**_γ_. As usual with the two H/C correlation peaks for **C**_γ_ are present, indicating that the γ-carbon bears two non-NMR-identical (diastereotopic) protons.

Supplemental Fig. S4 shows the additional assignment validation derived from the 3D NMR’s F1-F3 planes that represent ^1^H–^1^H TOCSY spectra at specific ^13^C NMR chemical shifts. Various planes from the β-ether **A**, phenylcoumaran **B**, and resinol **C** units illustrate the remarkable delineation of β-ether isomers achievable, consistently recognizing that the *threo* isomer possesses the lowest-δ proton.

### Lignin vs. polysaccharide acetylation

Non-cellulosic polysaccharides in the cell walls of plants, particularly dicots, are naturally *O*-acetylated (Donev et al., 2018). Specific xylan acetylation may facilitate hemicellulosic interactions with cellulose and lignin (Busse-Wicher et al., 2014), which is advantageous for certain industrial applications of plant biomass such as the production of xylooligosaccharides (XOS) (Wen et al., 2019), but it can also negatively impact the fermentation process (Pampulha et al., 1989). A recent report on the overexpression of the xylan acetylation gene (Zhang et al., 2024), known as the *Reduced Wall Acetylation* (*RWA*) gene, in poplar plants demonstrated an increase in lignin content and the lignin S/G ratio, along with enhanced xylan acetylation. However, this also resulted in reduced saccharification efficiency. Our NMR analysis of the extract-free cell wall residue revealed an absence of lignin acetylation in *RWA* poplars kindly provided by the aforementioned researchers (data not shown). Acetylation of lignin therefore represents a distinct aspect of cell wall acetylation.

### Mechanism of monolignol acetate incorporation into lignin

As elucidated by numerous studies conducted several decades ago (del Río et al., 2007; Lu et al., 2008), lignin acetylation is accomplished through the process of lignification using pre-formed monolignol acetate conjugates. The pathway from the monolignol to the monolignol acetate is catalyzed by an AMT enzyme (Fig. 3). Following export or transport to the cell wall’s lignifying zone, oxidation via an oxidase, laccase or peroxidase, produces the monolignol acetate phenolic radical. This radical is then ready for coupling with either another monolignol or monolignol acetate radical or, more commonly, with a phenolic radical on the growing lignin chain through endwise coupling (Freudenberg, 1965; Ralph et al., 2004b). The generation of the free radical at the phenolic terminus of the lignin oligomer/polymer remains slightly mysterious. It appears improbable that the immobile lignin entity can directly interact with the oxidase. Alternative pathways include radicalization from radical transfer from a mobile monolignol (or monolignol conjugate), or from another radical carrier (Ralph et al., 2009). Following the incorporation of the monolignol acetate into the growing polymer chain, further extension of the polymer can occur via the newly introduced phenolic endgroup.

**Figure 3.**
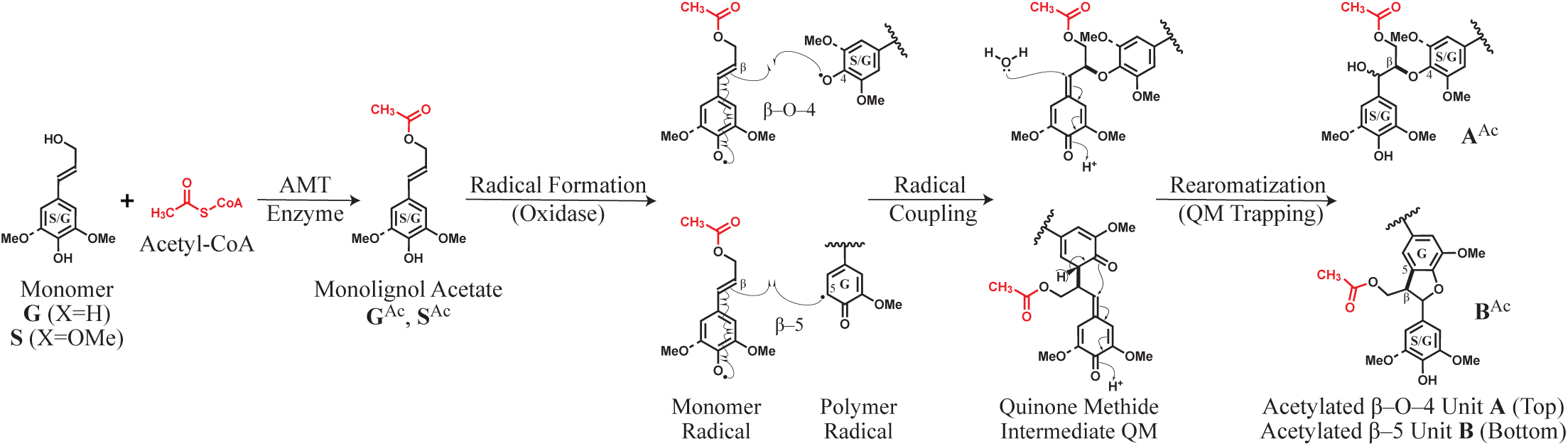
Monolignol acetate conjugate biosynthesis and its incorporation into lignin *via* radical coupling.

### Conclusions

Research on the mechanism of acylation and its impact on plant cell wall composition and overall plant growth remains limited. Our findings indicate that upregulation of the poplar monolignol acetylation gene *PtAMT1* resulted in substantial lignin acetylation, characterized by an increased percentage of G-type units in the lignin. The total lignin content remained unchanged. We conclude that *PtAMT1* is a poplar gene responsible for lignin acetylation, thereby addressing the previously unresolved issue of identifying a gene responsible for lignin acetylation.

In summary, our findings indicate that introducing acetates into lignin can effectively augment biomass acetate concentrations in an sustainable manner, without compromising polysaccharide content or the plant’s growth and development. Natural acetylation of lignin in biomass could potentially eliminate the requirement for mineral acid addition, commonly used in processes such as steam hydrolysis, thereby simplifying the refining process. Acetate can also serve as a sole carbon source for the production of valuable chemicals, including succinic acid, isobutanol, and isopropanol, through *Escherichia coli* fermentation (Gu et al., 2023; Narisetty et al., 2022; Yang et al., 2020).

## Materials and methods

### Screening and gene sequence analysis

To identify the acetyl-CoA:monolignol transferase candidate gene in *P. trichocarpa*, plant-specific BAHD acyl-CoA-dependent acyltransferases were selected based on the universally conserved motifs - HXXXD- and -DFGWG- (along with other variants such as -DFGFG-, -DFGWA-, -DFGWK-, - NFGWG-, -DLGFG-, -DYGWG-, and -NLGWG-) found in *P. trichocarpa* and other plants (D’Auria, 2006; Yu et al., 2009). These genes were retrieved from the NCBI GenBank database (https://www.ncbi.nlm.nih.gov) and Phytozome (https://phytozome-next.jgi.doe.gov). In total, 77 protein sequences were aligned with ClustalW2 v2.1 using default protein parameters (clustalw2 - INFILE=All_AMTs.fasta -TYPE=PROTEIN). The resulting multiple sequence alignment was used to infer an unrooted phylogeny under the neighbor-joining (NJ) criterion (Sato-Izawa et al., 2020). The Jones-Taylor-Thornton model (JTT model) and number of threads was set as 4 for the maximum likelihood method Pairwise evolutionary distances were computed from the MSA with Biopython v1.85 (Bio.Phylo.TreeConstruction.DistanceCalculator) using the BLOSUM62 distance model. To assess node support, we performed nonparametric bootstrap resampling with 1,000 replicates by resampling alignment columns with replacement. An NJ tree was reconstructed for each replicate and bootstrap support values were calculated as the percentage of replicates recovering the corresponding bipartition in the reference NJ tree (Stecher et al., 2020; Tamura et al., 2021). The final NJ tree was midpoint-rooted and internal node labels were limited to bootstrap percentages (all other internal labels suppressed). The tree was visualized using Matplotlib v3.9.4 via Biopython’s Phylo.draw; a scale bar is included to represent the displayed evolutionary distances (Supplemental Fig. S1B). Tree construction, bootstrapping, and visualization were performed in a local Python 3.11 environment (custom script available upon request). Well-known acyltransferase genes were eliminated based on the results of the sequence alignment and phylogenetic analysis. In the initial stage, three genes highly associated with shikimate *O*-hydroxycinnamoyltransferase in the NCBI database (www.ncbi.nlm.nih.gov) were selected as candidate genes for the acetyl-CoA:monolignol transferase (AMT). Among these three resulting hit sequences, one target sequence containing only the DFGWG motif was chosen. Subsequently, phylogenetic analysis of the selected candidate gene and all reduced-acetylation genes, including other identified BAHD transferase genes such as monolignol transferase genes (FMTs, PMTs) was reperformed using the maximum likelihood method. This method was complemented by comparison of translated amino acid sequences using MegaX software. Protein sequence identity analyses (Supplemental Table S1) were conducted using BLAST from the NCBI database. Conserved motifs of candidate genes were analyzed using the MEME (Multiple EM for Motif Elicitation) program (http://meme-suite.org/tools/meme), with random repetitions and optimum motif width set to 5–60. The predicted molecular mass of the PtAMT1 protein was calculated using Bioinformatics.org (http://www.bioinformatics.org/sms/prot_mw.htm).

### Sample preparation: Plant material, cell wall, and enzyme lignin

A chimeric poplar *PtAMT1* gene construct, controlled by the Arabidopsis *C4H* promoter, was introduced into hybrid poplar (*Populus tremula* × *Populus alba* clone INRA 717–1B4) using the Agrobacterium-mediated gene transformation method, as described by (Nabuqi et al., 2020), with minor modifications. Briefly, 1-month-old sterile poplar explants were used for Agrobacterium infection, and all gene transformation procedures were conducted on growth media solidified with Gelrite under sterile conditions. Basta (60 mg/L) was added to the culture medium for the selection and propagation of transgenic plants. The selected positive transgenic plants were subsequently analyzed by RT-PCR to confirm the expression of the transgenes (Supplemental Fig. S1A). Isolation of total RNA from poplar and profiling of *PtAMT1* gene transcripts in transgenic poplar were performed using the method previously described (Tamura et al., 2014; Nuoendagula et al., 2016). NEB Q5 High-Fidelity 2X Master Mix (Catalog Number M0492S) was utilized to amplify the full-length target DNA for 20 cycles, using cDNA from leaf tissue of the plant as the template. The elongation factor gene *EF1*β served as an internal standard. The primers for the target gene and the internal standard gene were as follows: *PtAMT1 gene* (forward: 5′-ATGGGTGCTACTGGAGGAGATG-3′; reverse: 5′-CTAGTTTGATGATGCAGATAGG-3′), *EF1*β gene (forward: 5′-AAGAGGACAAGAAGGCAGCA-3′; reverse: 5′-CTAACCGCCTTCTCCAACAC-3′). Three transgenic lines PtAMT-4, PtAMT-5, and PtAMT-18 were selected for further analysis. Wild-type poplar and all transgenic lines were vegetatively propagated on culture media in a growth chamber (22–25 °C; 16-h light: 8-h dark cycle). These plants were subsequently transplanted into soil in a greenhouse maintained at a temperature of 25 °C and a light-dark cycle of 16-h:8-h for further studies. A 6-month-old poplar stem was harvested for analysis of cell wall components after its bark removal. Extract-free cell wall residues, whole cell wall samples, and cellulase-enzyme-treated lignin were prepared using methods described previously (Lu et al., 2003; Wagner et al., 2007; Kim et al., 2010; Nabuqi et al., 2020).

### Lignin analysis

#### Thioglycolic acid lignin method

Lignin content was measured using the thioglycolic acid lignin method (Suzuki et al., 2009; Tsuyama et al., 2014). Briefly, 5-15 mg of cell wall residue in a 2 mL screw tube was suspended in 1 mL of 1 M hydrochloric acid (HCl) and 100 µL of thioglycolic acid (Sigma, USA), then incubated at 80 °C for 3 h. After centrifugation and washing with water, the pellet of cell wall residues was resuspended in 1 M NaOH solution and shaken at room temperature for 1 h. The supernatant (1 mL) collected after 10 min of centrifugation was used to reprecipitate the thioglycolic lignin by adding 3-4 drops of 6 M HCl. The resulting lignin pellet was dissolved in 1 M NaOH, and the lignin content was quantified using the calibration curve described by previously (Tsuyama et al., 2014), with absorbance measured at 280 nm.

#### Modified DFRC procedure

To assess the acetylation level of the predominant β-ether substructures (β– O–4 linkage) in lignin, the DFRC method, originally modified to the DFRC′ method for this purpose (Ralph et al., 1998b), was conducted using the labeled standards as previously described (Karlen et al., 2017; Regner et al., 2018). Briefly, extract-free ball-milled cell wall residue samples (20-50 mg) were stirred in 2-dram vials fitted with polytetrafluoroethylene pressure-release caps containing propionyl bromide in propionic acid (1:2 v/v, 5 mL). After stirring on a 50 °C sand bath for 16 h, the solvents were removed using a SpeedVac (Thermo Scientific SPD131DDA) with a run time of approximately 1.25 h, vacuum pressure of 1 torr, vacuum ramp 4, R/C lamp deactivated, temperature 50 °C. The crude films were then suspended in absolute ethanol (0.5 mL), dried under SpeedVac conditions (run time of approximately 20 min, other conditions as above), and subsequently suspended in a mixture of dioxane: propionic acid: water (5:4:1, v/v/v, 5 mL) along with nano-powdered zinc (250 mg). The vials were continually stirred in the dark at room temperature for 1 h (or sonicated for 1 h). Subsequently, the internal standard (300 μg/mL deuterated sinapyl alcohol diacetate) solution was added. The reaction mixtures were quantitatively transferred with dichloromethane (DCM; 10 mL) into separatory funnels charged with saturated ammonium chloride (10 mL) with additional DCM, 10 mL. Organic compounds were extracted with DCM (10 mL × 3), combined, dried over anhydrous sodium sulfate, filtered through filter paper, and the solvents were removed using rotary evaporation (water bath at 50 °C). Free hydroxyl groups on DFRC′ products were propionylated for 16 h in the dark using a solution of pyridine and propionic anhydride (1:1, v/v, 3 mL). The solvents were removed using a rotary evaporator (with a 60 °C water bath) to yield crude oily films. The products were transferred with GC-MS-grade DCM to GC-MS vials and analyzed using a triple-quadrupole GC/MS/MS instrument (Shimadzu GCMS-TQ8030) operating in MRM mode. MS and GC conditions are detailed in the Supplemental Materials.

#### 2D 1H–13C HSQC NMR

Six-month-old Poplar stem samples were utilized to prepare the enzyme lignin for analysis via ^1^H–^13^C 2D HSQC NMR spectroscopy. DMSO-d_6_/pyridine-d_5_ (4:1, v/v) was used as the solvent for samples run on a Bruker Biospin (Billerica, MA) Avance NEO 700 MHz spectrometer equipped with a 5-mm QCI ^1^H/^31^P/^13^C/^15^N cryoprobe with inverse geometry (proton coils closest to the sample), as described previously (Kim et al., 2008; Kim et al., 2010). All NMR experiments used Bruker’s standard pulse programs. An adiabatic ^1^H–^13^C 2D HSQC experiment (hsqcetgpsisp2.2; phase-sensitive gradient-edited 2D HSQC using adiabatic pulse sequences for inversion and refocusing) was used to collect the main data (Kupče et al., 2007). HSQC experiments were conducted with the following parameters: acquired from 11.66 to −0.66 ppm in F2 (^1^H) with 3448 datapoints (acquisition time, 200 ms) and 215 to −5 ppm in F1 (^13^C) with 618 increments (F1 acquisition time, 8 ms) of 24 scans with a 1 s interscan delay; the d24 delay was set to 0.89 ms (⅛_J_, J = 145 Hz). The total acquisition time for a sample was 5 h 22 min. Processing used typical matched Gaussian apodization (LB = −0.5, GB = 0.001) in F2 and squared cosine-bell in F1 (without linear prediction). Spectra were referenced (as customary) using the central DMSO solvent peak in the HSQC spectra (δ_C_ 39.5, δ_H_ 2.49 ppm); these shifts are offset compared to externally-referenced spectra in which case the carbon shifts should be shifted by -0.276 ppm and the proton shifts by +0.0011 ppm. Volume-integration of contours in HSQC plots was performed using TopSpin 4.4 (Mac) software, without the use of correction factors.

#### 3D ^1^H–^13^C TOCSY-HSQC NMR

The 3D spectrum acquired for the 2D partial planes shown in Supplemental Figs. S3 and S4 was acquired on the same 700 MHz spectrometer and using the same enzyme lignins (EL) as described above. The 3D gradient-selected TOCSY-HSQC experiment was Bruker’s standard mlevhsqcetgp3d experiment acquiring just the oxygenated-aliphatic region using the following parameters: 952 × 128 × 64 datapoints taken over sweep widths of 3.4 (^1^H, F3, acquisition time 200 ms), 59 (^13^C, F2, acquisition time 6.16 ms), and 3.4 (^1^H, F1, acquisition time 13.45 ms) ppm centered at 4.25, 75.5, and 4.25 ppm, with 16 scans per increment, a TOCSY mixing time d9 of 60 ms, a d4 delay of 1.78 ms (¼_J_, J = 140 Hz), and a relaxation delay time d1 of 1 s. The total experiment time was 46 h. Processing used matched Gaussian (LB = −0.2, GB = 0.001) apodization in F3, cosine-bell-squared (qsine, SSB =2) in F2 and F1], and processed to 1k × 512 × 512 datapoints with forward linear prediction (32 coefficients) in F2 and F1. Similar spectra can be found in various of our references (Ralph et al., 1999; Ralph et al., 2004a; Lu et al., 2015; del Río et al., 2017; Tobimatsu et al., 2019).

## Acknowledgements

We extend our sincere gratitude to the National Institute for Agricultural Research, Food and Environment (INRAE), France, for providing the INRA 717-1B4 clone (*P. tremula* × *P. alba*) under a Material Transfer Agreement (MTA). Our heartfelt thanks also go to Professor Kyung Hwan Han from the Department of Horticulture at Michigan State University for generously supplying poplar tissue culture materials, and to Professor Chung-Jui (CJ) Tsai from the Department of Genetics at the University of Georgia for her invaluable assistance with the poplar gene transformation methods employed in this study. We also acknowledge Dr. Aymerick Eudes from Lawrence Berkeley National Laboratory, Joint BioEnergy Institute, for providing the wood powder from transgenic *Arabidopsis* harboring the *CFAT* gene from *Petunia hybrida*. We express our appreciation to Dr. Steve Karlen, Matt Regner, and Vitaliy I. Timokhin from the Great Lakes Bioenergy Research Center, University of Wisconsin-Madison, for providing DFRC standard compounds and offering valuable advice on GC-MS analysis. Sally Ralph, US Forest Products Laboratory, is gratefully acknowledged for providing precursors to the β-ether models. Additionally, we thank Dr. Jin-Gui Chen and Tao Yao from Oak Ridge National Laboratory/University of Georgia for kindly supplying the wood powder from transgenic poplar overexpressing the *Reduced Wall Acetylation* (*RWA*) gene. We thank Prof. Shawn Mansfield for his advice on analysis, suggestions for this project, and his continued interest. Professor Clint Chapple kindly provided the empty plant expression vector utilized in this study.

