## Supplementary material for "A Newly Identified Gene Controls Lignin Acetylation in Poplar": Figures

### Discovery and validation of a gene for lignin acetylation in Poplar

#### TABLE OF CONTENTS

**GC-MS parameters for DFRC′ analyses**
**NMR spectra**
**Supplemental Figure S1.** Evolutionary relationship of PtAMT and related proteins, and RT-PCR.
**Supplemental Figure S2.** HSQC plot as in Fig. 2A but showing synthesized model data.
**Supplemental Figure S3.** 2D HSQC planes from a 3D TOCSY-HSQC spectrum.
**Supplemental Figure S4.** 2D TOCSY slices from the 3D TOCSY-HSQC experiment.
**Supplemental Figure S5.** Aromatic regions of the 2D HSQC NMR spectra of enzyme lignins.
**Table S1.** Protein sequence identity between PtAMTa enzymes and related enzymes.

#### GC-MS parameters for DFRC′ analyses

GDFRC samples were analyzed using a triple-quadrupole GC/MS/MS device (Shimadzu GCMS-TQ8030) operating in MRM mode.

***Gas Chromatography***
Gas Chromatograph: GC-2010 Plus
Column: RXi-5Sil MS 30 m × 0.25 mm × 0.25 µm (Restek 13623)
Column Oven Temp: 150.0 °C
Inlet: 250 °C
Split liner with glass wool (Shimadzu 220-90784-00)
Split injection (20:1)
Column Flow: 0.70 mL/min
Linear Velocity: 44.9 cm/s
Purge Flow: 1.0 mL/min
Total Flow: 15.7 mL/min
High Pressure Injection: off
Carrier Gas: Helium
Carrier Gas Saver: off
Splitter Hold: off,
Oven Program: 150 °C, hold 1 min, ramp 10 °C/min to 300 °C, hold 29.0 min

***Mass Spectrometry***
Mass Spectrometer: GCMS-TQ8030
MS interface 300 °C
Analysis time 45.0 min
Ion Source: 275 °C
Mode: Electron impact (EI), 70 eV
Interface Temp: 300 °C
Solvent Cut Time: 4.00 min
Detector Gain Mode: Relative to the Tuning Result
Detector Gain: 1.06 kV +0.00 kV
Threshold: 0
Acquire Data without Using CID Gas(Q3Scan): OFF
Operation Mode: Multiple Reaction Monitoring (MRM)
Gas: Argon, 200 kPa
Q1 resolution: 0.8 u (unit)
Q3 resolution: 0.8 u (unit)
Detector: Electron multiplier 0.97 kV.

***MS Table details***
Group1 – Event 1: Compound Name: Q3Scan Start Time: 4.20 min, End Time: 45.0 min, Acq. Mode: Q3 Scan, Event Time: 0.150 s, Scan Speed:5000, Start m/z: 100, End m/z: 800, Q1 Resolution: -, Q3 Resolution: -; Group1 – Event 2: Compound Name: HOPrOAc, Start Time: 4.20min, End Time: 45.00min, Acq. Mode: MRM, Event Time: 0.050 s, Q1 Resolution: Unit, Q3 Resolution: Unit, Ch1-m/z: (Precursor) 248.00, Ch2-m/z: (Precursor) 248.00, Ch3-m/z: (Precursor) 248.00; Group1 – Event 3: Compound Name: H-DOPr, Start Time: 4.20 min, End Time: 45.00 min, Acq. Mode: MRM, Event Time: 0.050 s, Q1 Resolution: Unit, Q3 Resolution: Unit, Ch1-m/z: (Precursor) 262.00, Ch2-m/z: (Precursor) 262.00, Ch3-m/z: (Precursor) 262.00; Group1 - Event 4: Compound Name: G-OPrOAc, Start Time: 4.20 min, End Time: 45.00 min, Acq. Mode: MRM, Event Time: 0.050 s, Q1 Resolution: Unit, Q3 Resolution: Unit, Ch1-m/z: (Precursor) 278.00, Ch2-m/z: (Precursor) 278.00, Ch3-m/z: (Precursor) 278.00; Group1 – Event 5: Compound Name: G-DPr, Start Time: 4.2 min, End Time: 45 min, Acq. Mode: MRM, Event Time: 0.050 s, Q1 Resolution: Unit, Q3 Resolution: Unit, Ch1-m/z: (Precursor) 292.00, Ch2-m/z: (Precursor) 292.00, Ch3-m/z: (Precursor) 292.00; Group1 – Event 6: Compound Name: S-OPrOAc, Start Time: 4.20 min, End Time: 45.00 min, Acq. Mode: MRM, Event Time: 0.050 s, Q1 Resolution: Unit, Q3 Resolution: Unit,

Ch1-m/z: (Precursor) 308.00, Ch2-m/z: (Precursor) 308.00, Ch3-m/z: (Precursor) 308.00, Ch4-m/z: (Precursor) 294.00; Group1 – Event 7: Compound Name: S-DOPr, Start Time: 4.20 min, End Time: 45.00 min, Acq. Mode: MRM, Event Time: 0.050 s, Q1 Resolution: Unit, Q3 Resolution: Unit, Ch1-m/z: (Precursor) 322.00, Ch2-m/z: (Precursor) 322.00, Ch3-m/z: (Precursor) 322.00, Ch4-m/z: (Precursor) 294.00; Sample Inlet Unit: GC; [MS Program]; Use MS Program: OFF.

**NMR spectra**
All spectra were acquired as described the text's Materials and Methods Section. All raw NMR data, Bruker TopSpin format, may be obtained from the authors upon request. All the data associated with the authentic model compounds (Fig. S2) will be deposited in this year's release of our NMR Database (Ralph et al., 2025).

**A**

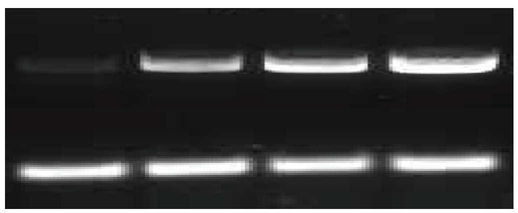


**B**

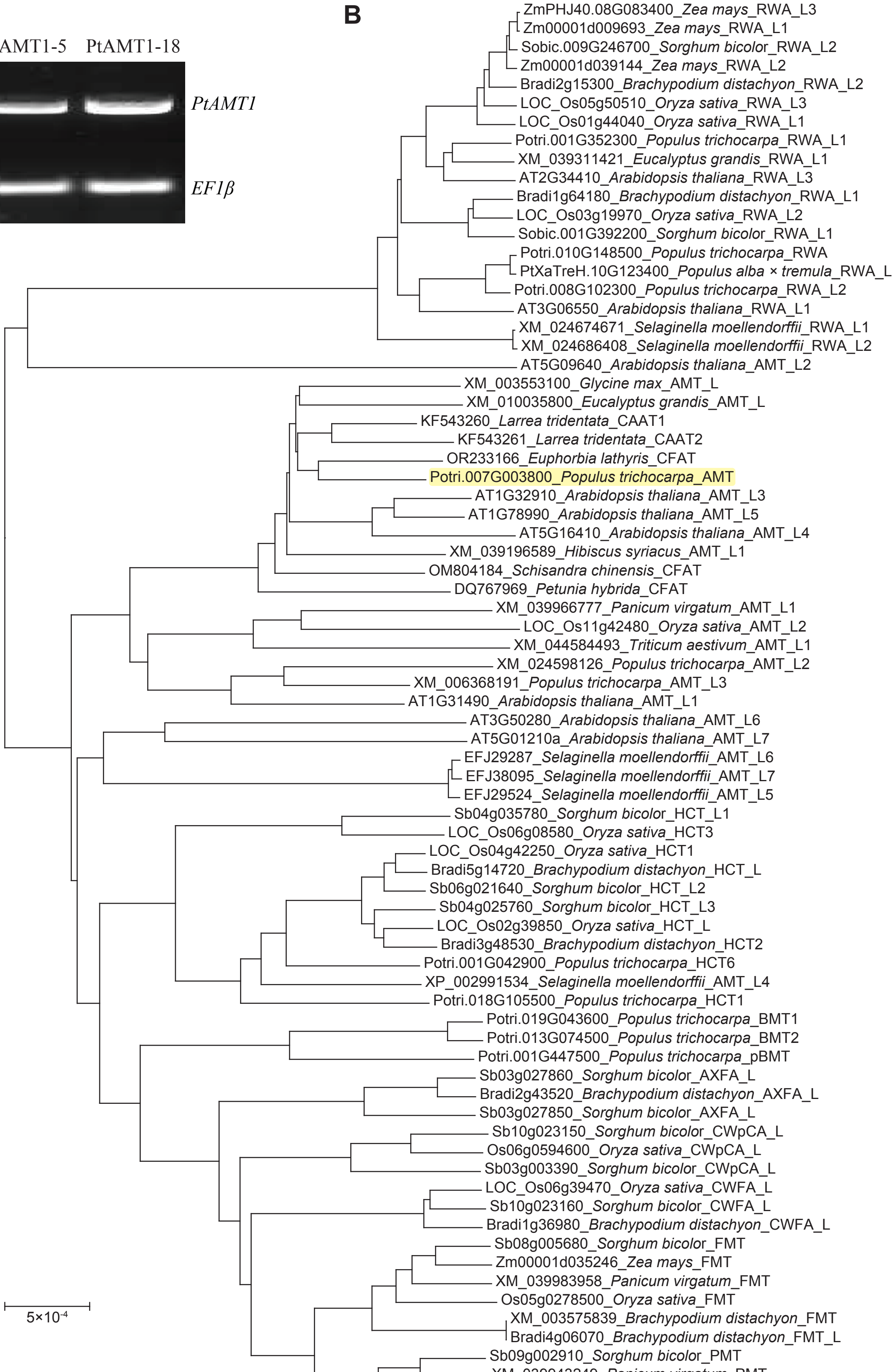


**Supplemental Figure S1. A)** Reverse transcription-polymerase chain reaction (RT-PCR) analysis of gene of interest transcripts in leaves of WT and transgenic lines, with EF1β (elongation factor 1β) transcripts analyzed as a control. **B)** Peptide sequence evolutionary relationship of PtAMT and related proteins. Abbreviations: FMT: feruloyl-CoA:monolignol transferase; PMT: *p*-coumaroyl-CoA:monolignol transferase; CW*p*CA: *p*-coumaroylated cell wall; CWFA: feruloylated cell wall (primarily on arabinoxylan). HCT: hydroxycinnamoyl-CoA:shikimate transferase; CAAT: cinnamyl alcohol:acyl-CoA transferase (acyl-CoA:cinnamyl alcohol transferase); CFAT: coniferyl alcohol:acetyl-CoA transferase (acetyl-CoA:coniferyl alcohol transferase), also a generic CAAT; AMT: acetyl-CoA:monolignol transferase; AXFA: arabinoxylan:ferulic acid (or feruloyl-CoA) transferase; RWA: reduced wall acetylation; L: Like.

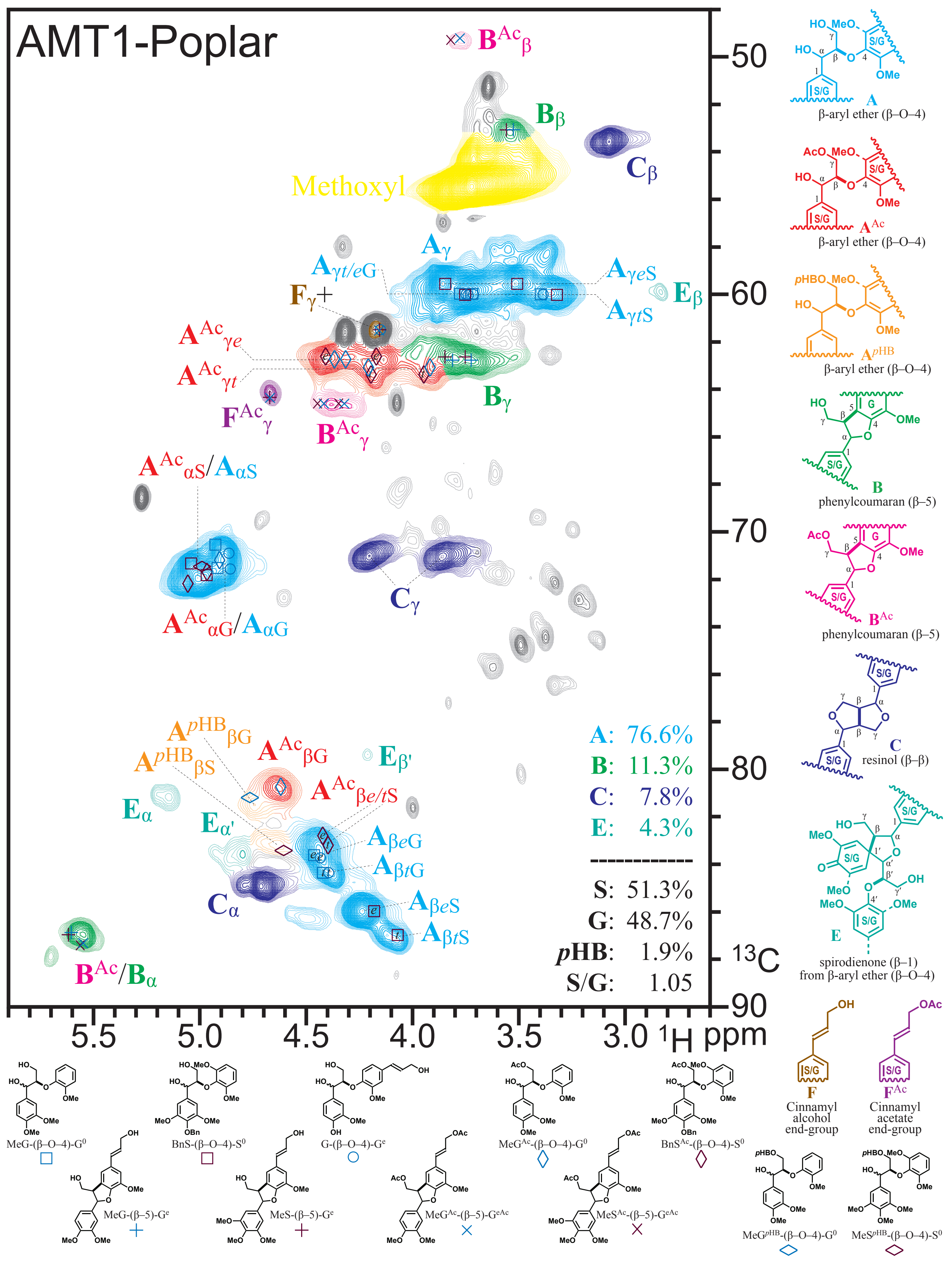
AMT1-Poplar
Methoxyl
A: 76.6%
B: 11.3%
C: 7.8%
E: 4.3%
S: 51.3%
G: 48.7%
pHB: 1.9%
S/G: 1.05
¹H ppm
¹³C
A
β-aryl ether (β–O–4)
A^Ac
β-aryl ether (β–O–4)
A^pHB
β-aryl ether (β–O–4)
B
phenylcoumaran (β–5)
B^Ac
phenylcoumaran (β–5)
C
resinol (β–β)
E
spirodienone (β–1)
from β-aryl ether (β–O–4)
F
Cinnamyl alcohol end-group
F^Ac
Cinnamyl acetate end-group
MeG-(β–O–4)-G⁰
BnS-(β–O–4)-S⁰
G-(β–O–4)-Gᵉ
MeG^Ac-(β–O–4)-G⁰
BnS^Ac-(β–O–4)-S⁰
MeG-(β–5)-Gᵉ
MeS-(β–5)-Gᵉ
MeG^Ac-(β–5)-G^eAc
MeS^Ac-(β–5)-G^eAc
MeG^pHB-(β–O–4)-G⁰
MeS^pHB-(β–O–4)-S⁰

*Previous page*

**Supplemental Figure S2.** The same HSQC plot as in Fig. 2A in the main paper but showing chemical shift markers for the synthesized models at the bottom, instructively with (× and ◊) and without (+, □, and ○) γ-acetylation, and (sideways-◊) for *p*-hydroxybenzoylation.

*Next page*

**Supplemental Figure S3.** $^{1}H$–$^{13}C$ 2D HSQC (F2-F3) planes from the 3D TOCSY-HSQC spectrum. Selected F2-F3 planes were extracted for figures B-N at the noted proton chemical shift in F1. The normal 2D HSQC spectrum from **A)** is lightly displayed underneath each plot **B)-N)** for reference; relevant contours are colored as in **A)** and Fig. 2A in the main paper, whereas other/breakthrough contours are simply colored a darker gray than the gray used in the underlying plot, i.e., we have not edited these spectra other than to color the contours of interest. The power of these spectra is that they show all sidechain peaks related to a given structure and provide proof, for example, that the peak labeled $\mathbf{B}^{Ac}{}_{\beta}$ in **B)** and **C)** is indeed from the same component as that labeled $\mathbf{B}^{Ac}{}_{\gamma}$, and that the $\mathbf{B}^{Ac}{}_{\alpha}$ peak is not resolved from the normal $\mathbf{B}_{\alpha}$ peak.

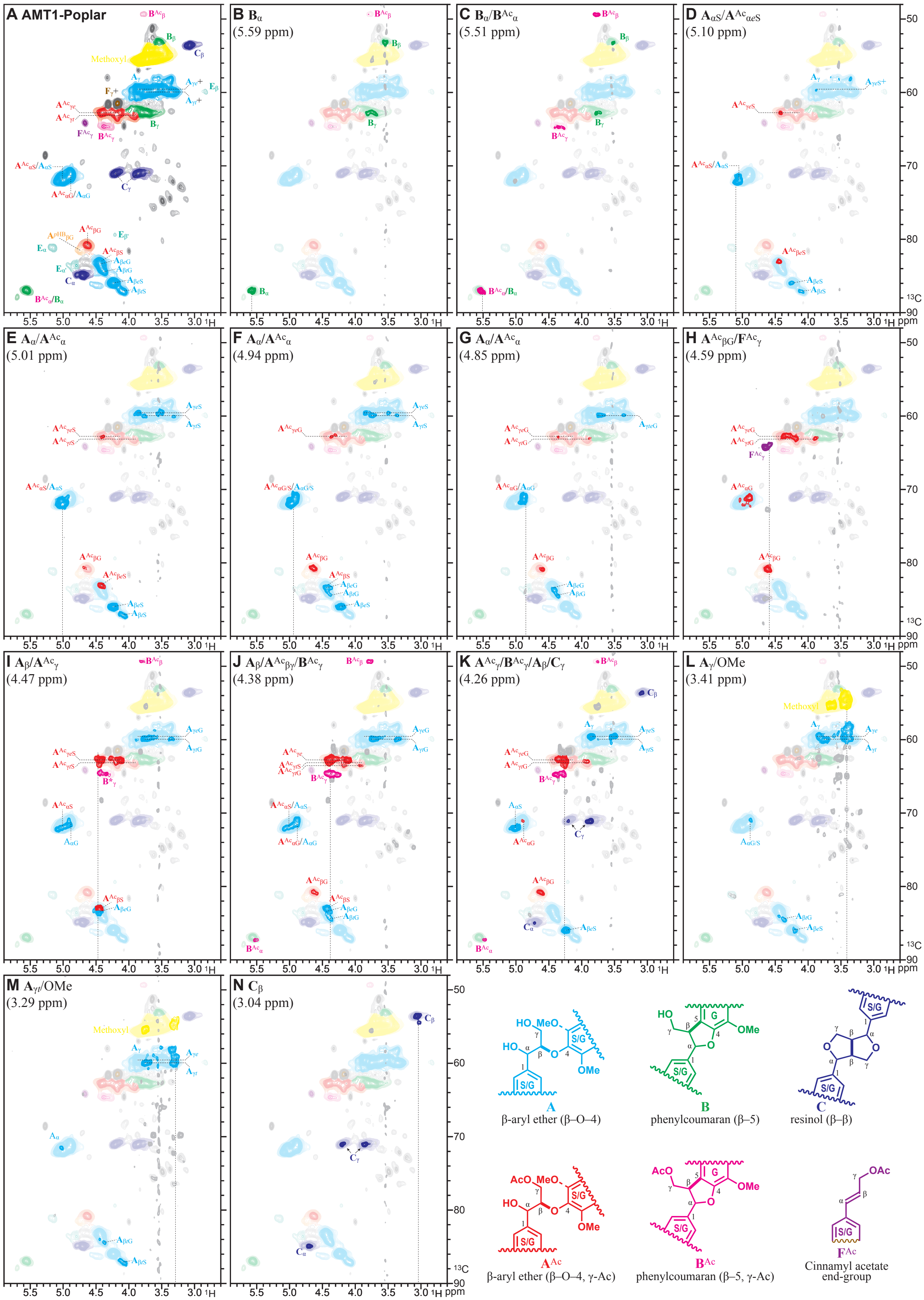

A AMT1-Poplar
B Bα (5.59 ppm)
C Bα/BAcα (5.51 ppm)
D AαS/AAcαeS (5.10 ppm)
E Aα/AAcα (5.01 ppm)
F Aα/AAcα (4.94 ppm)
G Aα/AAcα (4.85 ppm)
H AAcβG/FAcγ (4.59 ppm)
I Aβ/AAcγ (4.47 ppm)
J Aβ/AAcβγ/BAcγ (4.38 ppm)
K AAcγ/BAcγ/Aβ/Cγ (4.26 ppm)
L Aγ/OMe (3.41 ppm)
M Aγ/OMe (3.29 ppm)
N Cβ (3.04 ppm)
Methoxyl
A
β-aryl ether (β–O–4)
B
phenylcoumaran (β–5)
C
resinol (β–β)
AAc
β-aryl ether (β–O–4, γ-Ac)
BAc
phenylcoumaran (β–5, γ-Ac)
FAc
Cinnamyl acetate end-group

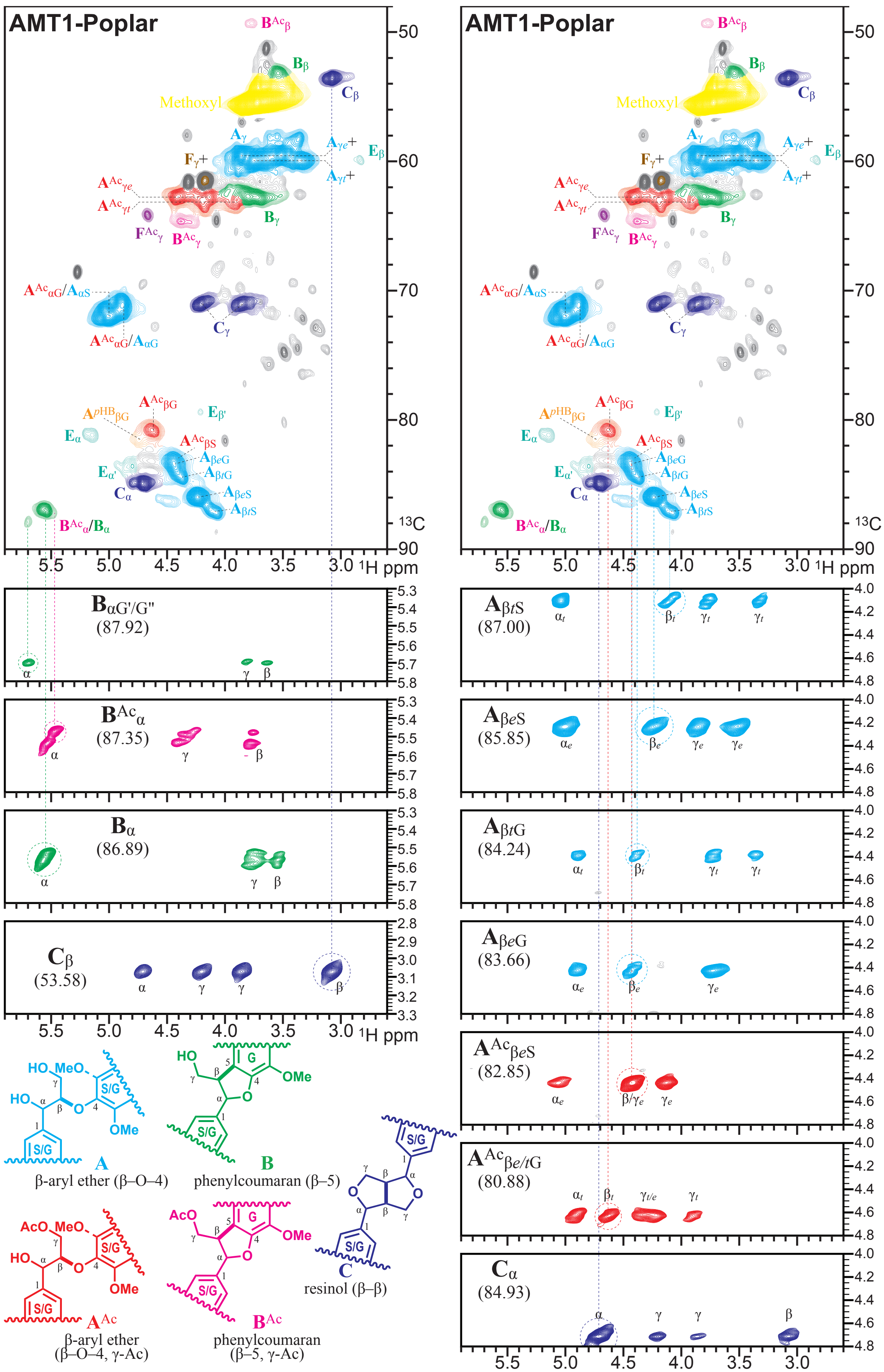


**Supplemental Figure S4.** 2D TOCSY (F1–F3) planes from the 3D TOCSY-HSQC experiment over a small proton chemical shift range in the F3 dimension are shown under (and aligned with) the same 2D HSQC spectra as those from Fig. 2A in the main paper and Supplemental Fig. S2 here. The easily-assigned resinol **C** correlations are shown from the $C_{\alpha}$ and $C_{\beta}$ carbon planes, and the various normal (non-γ-acetylated) and γ-acetylated phenylcoumaran **B** and β-ether **A** peaks from various carbon planes are shown. These particularly clean partial-spectra often allow beautiful distinction between *threo* (*t*) and *erythro* (*e*) isomers, and between $A_{\beta S}$ *vs* $A_{\beta G}$ moieties in the polymer, resolving peaks that are often overlapped in the 2D HSQC spectrum.

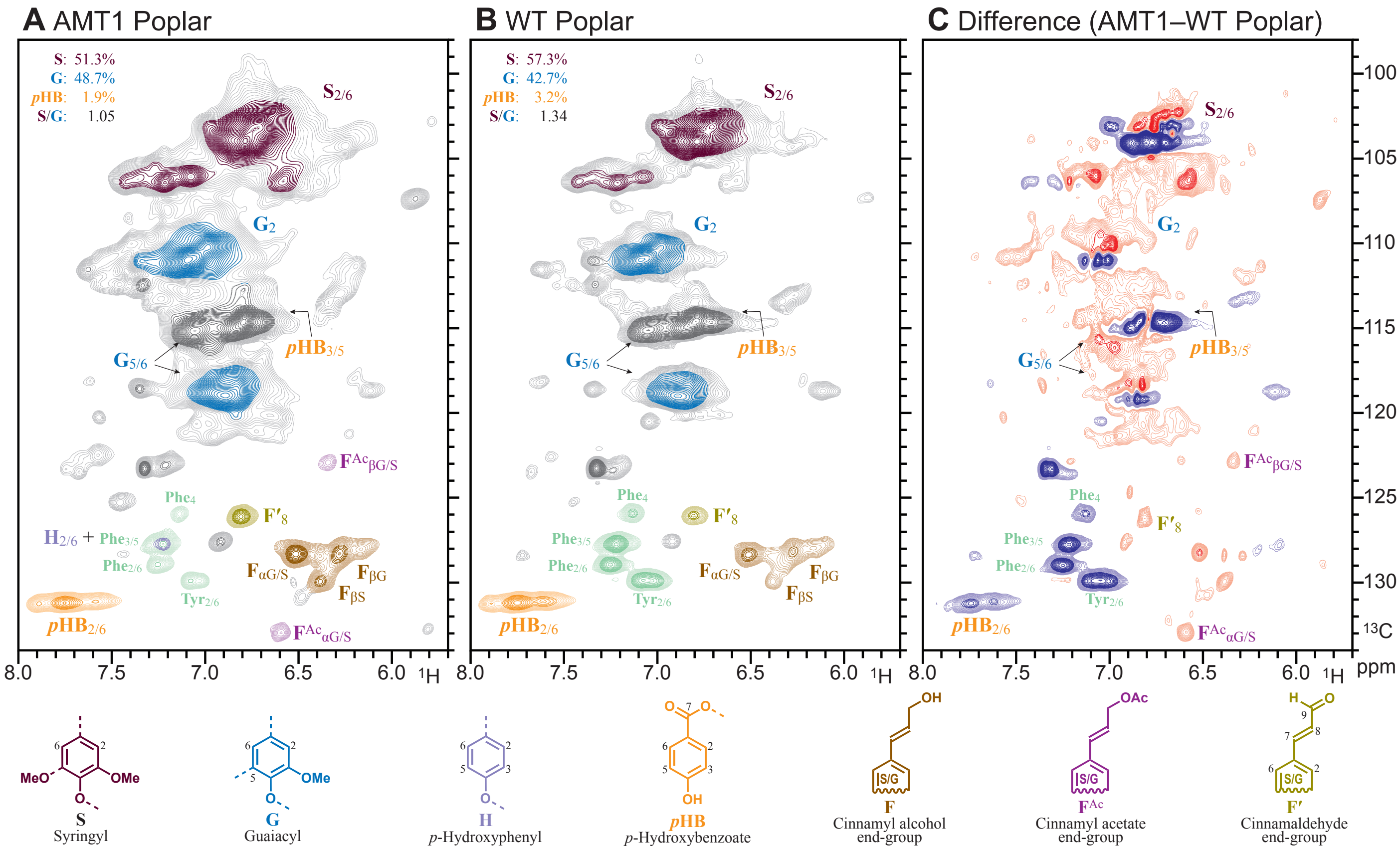


**Supplemental Figure S5.** Aromatic (and double-bond) regions of the 2D HSQC NMR spectra of enzyme lignins (EL). **A)** The AMT1 transgenic with the acetyl-CoA:monolignol transferase expressed. **B)** The WT control poplar. **C)** The 2D difference spectrum (AMT1 minus WT) using reasonable nulling of the guaiacyl **G**$_2$ peak, revealing the compositional differences. The warm colors represent components elevated in the transgenic line *vs* WT, whereas cold colors represent depleted components relative to the guaiacyl peaks. As can be discerned, peaks from *p*-hydroxybenzoates (***p*HB**) in particular were markedly lower, and the **S**-level is lower and has more diverse types. The cinnamyl acetate end-group (**F**$_{Ac}$) levels are revealed to logically be at higher levels given that they are essentially not present in the WT. For whatever reason, the level of residual proteins (as seen by the **Phe** and **Tyr** correlations) are also lower in the AMT1 lignin. The lighter underlaid plots are at a 4× lower contour level. Relative levels of **S**, **G**, and ***p*HB** components tabulated in the upper-left corner are from uncorrected volume-integrals.

**Table S1.** Protein sequence identity between PtAMTa enzymes and the identified lignin pathway enzymes, monolignol transferases, and reduced wall acetylation (RWA) enzymes.

| Name (Gene ID, Species and Gene name) | Max Score | Total Score | Query Cover | Identity (%) | AA length |
|---|---|---|---|---|---|
| Potri.007G003800_*Populus Trichocarpa* _AMT | 1018 | 1018 | 100% | 100.00% | 492 |
| KF543260_*Larrea Tridentata* _CAAT1 | 682 | 682 | 97% | 69.87% | 463 |
| OR233166_*Euphorbia Lathyris* _CFAT | 625 | 625 | 96% | 67.30% | 457 |
| XM_039196589_*Hibiscus Syriacus* _AMT_Like1 | 608 | 608 | 96% | 62.74% | 455 |
| KF543261_*Larrea Tridentata* _CAAT2 | 589 | 589 | 96% | 64.12% | 463 |
| XM_010035800_*Eucalyptus Grandis* _AMT_Like | 588 | 588 | 99% | 61.86% | 487 |
| XM_003553100_*Glycine Max* _AMT_Like | 553 | 553 | 98% | 61.29% | 472 |
| OM804184_*Schisandra Chinensis* _CFAT | 550 | 550 | 97% | 61.01% | 455 |
| AT1G78990_*Arabidopsis Thaliana* _AMT_Like5 | 549 | 549 | 98% | 60.37% | 455 |
| AT1G32910_*Arabidopsis Thaliana* _AMT_Like3 | 549 | 549 | 96% | 60.62% | 464 |
| AT5G16410_*Arabidopsis Thaliana* _AMT_Like4 | 490 | 490 | 97% | 52.58% | 481 |
| DQ767969_*Petunia Hybrida* _CFAT | 488 | 488 | 96% | 56.12% | 454 |
| XM_006368191_*Populus Trichocarpa* _AMT_Like3 | 282 | 282 | 96% | 36.97% | 448 |
| AT1G31490_*Arabidopsis Thaliana* _AMT_Like1 | 266 | 266 | 96% | 34.17% | 444 |
| XM_039966777_*Panicum Virgatum* _AMT_Like1 | 158 | 158 | 96% | 27.50% | 467 |
| Potri.018G105500_*Populus Trichocarpa* _HCT1 | 135 | 135 | 79% | 27.50% | 430 |
| XM_044584493_*Triticum Aestivum* _AMT_Like1 | 130 | 130 | 96% | 27.31% | 477 |
| Potri.001G042900_*Populus Trichocarpa* _HCT6 | 127 | 127 | 88% | 25.96% | 431 |
| LOC_OS11g42480_*Oryza Sativa* _AMT_Like2 | 126 | 126 | 95% | 24.31% | 442 |
| XP_002991534_*Selaginella Moellendorffii* _AMT_Like4 | 123 | 123 | 84% | 26.42% | 449 |
| SB06g021640_*Sorghum Bicolor* _HCT_Like2 | 118 | 118 | 95% | 25.87% | 441 |
| Bradi3g48530_*Brachypodium Distachyon* _HCT2 | 114 | 114 | 78% | 26.72% | 442 |
| SB04g035780_*Sorghum Bicolor* _HCT_Like1 | 112 | 112 | 90% | 26.74% | 451 |
| EFJ29524_*Selaginella Moellendorffii* _AMT_Like5 | 112 | 112 | 91% | 26.41% | 444 |
| EFJ29287_*Selaginella Moellendorffii* _AMT_Like6 | 111 | 111 | 91% | 26.41% | 444 |
| LOC_OS02g39850_*Oryza Sativa* _HCT_Like | 110 | 110 | 88% | 24.01% | 442 |
| EFJ38095_*Selaginella Moellendorffii* _AMT_Like7 | 109 | 109 | 91% | 25.80% | 444 |
| LOC_OS04g42250_*Oryza Sativa* _HCT1 | 108 | 108 | 84% | 25.34% | 442 |
| Potri.001G447500_*Populus Trichocarpa* _pBMT | 106 | 106 | 83% | 25.41% | 465 |
| SB04g025760_*Sorghum Bicolor* _HCT_L3 | 103 | 103 | 88% | 24.78% | 448 |
| Potri.019G043600_*Populus Trichocarpa* _BMT1 | 102 | 102 | 75% | 25.19% | 460 |
| Potri.013G074500_*Populus Trichocarpa* _BMT2 | 99 | 99 | 80% | 24.24% | 459 |
| LOC_OS06g08580_*Oryza Sativa* _HCT3 | 97.8 | 97.8 | 84% | 24.94% | 445 |
| Bradi5g14720_*Brachypodium Distachyon* _HCT_Like | 92.8 | 92.8 | 79% | 24.32% | 443 |
| AT3G50280_*Arabidopsis Thaliana* _AMT_Like6 | 78.2 | 78.2 | 78% | 21.81% | 443 |
| SB10g023150_*Sorghum Bicolor* _CWpCA_Like | 75.1 | 75.1 | 25% | 35.11% | 446 |
| XM_024598126_*Populus Trichocarpa* _AMT_Like2 | 72.4 | 72.4 | 43% | 26.85% | 274 |
| SB03g027850_*Sorghum Bicolor* _AXFA_Like | 72.4 | 72.4 | 41% | 30.14% | 428 |
| SB03g027860_*Sorghum Bicolor* _AXFA_Like | 70.9 | 70.9 | 37% | 30.89% | 426 |
| Bradi2g43520_*Brachypodium Distachyon* _AXFA_Like | 70.5 | 70.5 | 26% | 34.78% | 423 |
| OS06g0594600_*Oryza Sativa* _CWpCA_Like | 70.1 | 70.1 | 25% | 33.33% | 443 |
| AT5G01210a_*Arabidopsis Thaliana* _AMT_Like7 | 68.9 | 68.9 | 77% | 24.94% | 475 |
| SB10g023160_*Sorghum Bicolor* _CWFA_Like | 67.4 | 67.4 | 40% | 30.23% | 438 |
| Bradi1g36980_*Brachypodium Distachyon* _CWFA_Like | 66.2 | 66.2 | 36% | 30.05% | 432 |
| OS01g0291500_*Oryza Sativa* _PMT1 | 63.5 | 63.5 | 37% | 31.44% | 439 |
| LOC_OS01g18744_*Oryza Sativa* _PMT_Like | 63.5 | 63.5 | 37% | 31.44% | 439 |
| Bradi2g36910_*Brachypodium Distachyon* _PMT | 63.2 | 63.2 | 24% | 34.13% | 449 |
| SB03g003390_*Sorghum Bicolor* _CWpCA_Like | 60.1 | 60.1 | 27% | 30.56% | 417 |
| SB09g002910_*Sorghum Bicolor* _PMT | 60.1 | 60.1 | 23% | 34.45% | 436 |
| XM_003575839_*Brachypodium Distachyon* _FMT | 59.3 | 59.3 | 28% | 31.76% | 442 |
| LOC_OS06g39470_*Oryza Sativa* _CWFA_Like | 57.4 | 57.4 | 24% | 31.06% | 431 |
| OS05g0278500_*Oryza Sativa* _FMT | 57.4 | 57.4 | 27% | 32.39% | 432 |
| XM_039943249_*Panicum Virgatum* _PMT | 57.4 | 57.4 | 46% | 25.74% | 427 |
| SB08g005680_*Sorghum Bicolor* _FMT | 56.6 | 56.6 | 26% | 33.33% | 440 |
| XM_039983958_*Panicum Virgatum* _FMT | 54.7 | 54.7 | 26% | 34.75% | 436 |
| ZM00001d035246_*Zea Mays* _FMT | 53.1 | 53.1 | 26% | 33.33% | 433 |
| Bradi4g06070_*Brachypodium Distachyon* _FMT_Like | 51.2 | 51.2 | 28% | 28.93% | 453 |
